# Hierarchical cortical transcriptome disorganization in autism

**DOI:** 10.1101/042937

**Authors:** Michael V. Lombardo, Eric Courchesne, Nathan E. Lewis, Tiziano Pramparo

**Author notes:** **Corresponding Author**: Michael V. Lombardo.

## Abstract

**Background:** Autism spectrum disorders (ASD) are etiologically heterogeneous and complex. Functional genomics work has begun to identify a diverse array of dysregulated transcriptomic programs (e.g., synaptic, immune, cell cycle, DNA damage, WNT signaling, cortical patterning and differentiation) potentially involved in ASD brain abnormalities during childhood and adulthood. However, it remains unclear whether such diverse dysregulated pathways are independent of each other or instead reflect coordinated hierarchical systems-level pathology.

**Methods:** Two ASD cortical transcriptome datasets were re-analyzed using consensus weighted gene co-expression network analysis (WGCNA) to identify common coexpression modules across datasets. Linear mixed-effect models and Bayesian replication statistics were used to identify replicable differentially expressed modules. Eigengene network analysis was then utilized to identify between-group differences in how co-expression modules interact and cluster into hierarchical meta-modular organization. Protein-protein interaction analyses were also used to determine whether dysregulated co-expression modules show enhanced interactions.

**Results:** We find replicable evidence for 10 gene co-expression modules that are differentially expressed in ASD cortex. Rather than being independent non-interacting sources of pathology, these dysregulated co-expression modules work in synergy and physically interact at the protein level. These systems-level transcriptional signals are characterized by downregulation of synaptic processes coordinated with upregulation of immune/inflammation, response to other organism, catabolism, viral processes, translation, protein targeting and localization, cell proliferation, and vasculature development. Hierarchical organization of meta-modules (clusters of highly correlated modules) is also highly affected in ASD.

**Conclusions:** These findings highlight that dysregulation of the ASD cortical transcriptome is characterized by the dysregulation of multiple coordinated transcriptional programs producing synergistic systems-level effects that cannot be fully appreciated by studying the individual component biological processes in isolation.

---

The pathophysiology behind atypical brain development in autism spectrum disorders (ASD) is highly complex. Elegant biological studies are continually unveiling an ever more diverse array of etiological factors and neurodevelopmental processes associated with ASD (e.g., [1–17]). With such diversity, key questions arise as to what are the consistent and robust mechanisms involved and whether such mechanisms point to many independent disrupted pathways or some convergence on a few common pathways affecting large-scale biological systems and/or interactions between such systems [18, 19]. One way to test this question is to examine pathophysiology at a level above genetics and non-genetic perturbations, such as the transcriptome, and examine whether the diversity of disrupted transcriptomic signals converge onto many independent or interacting systems.

Seminal work examining cortical transcriptome dysregulation in ASD has highlighted the dysregulation of multiple transcriptional programs. These programs include cell cycle/DNA damage, WNT signaling, cortical patterning and differentiation, and immune/inflammation at young ASD ages [5] and apoptosis, repair and remodeling, synaptic and immune/inflammation functions at older ages [8, 20, 21]. However, it remains unclear if such pathways are independently dysregulated or whether there is synergy between multiple dysregulated pathways. For example, prior work has shown downregulated synaptic and upregulated immune/inflammation signals in ASD cortical tissue [8, 20, 21]. Pointing towards the idea that such dysregulated signals may not be independent, strong correlations are found between these dysregulated modules when collapsing data across both groups [8, 20]. While this observation is important for generally suggesting statistical dependency between modules, more evidence is needed to suggest that such potentially synergistic effects among interacting modules are dysregulated in ASD. To test the hypothesis that transcriptome dysregulation in ASD extends beyond the level of single dysregulated co-expression modules and involves dysregulation spanning interactions between larger systems-level processes, differences in between-module correlations need to be investigated. Furthermore, tests should also go beyond observations of statistical dependency in co-expression and test whether there are enhanced direct physical interactions between the protein products of such dysregulated modules compared to unaffected modules. An enhancement of direct physical protein interactions amongst dysregulated versus non-dysregulated modules would further suggest plausibility for synergistic interactions across disparate biological processes conferred by each individual co-expression module.

We examined this topic via re-analysis of two existing datasets that investigated multiple cortical regions. We tested the hypothesis that diverse molecular mechanisms are hierarchically disrupted in the cortical transcriptome of ASD and reflect interacting systems-level pathology rather than multiple independent types of molecular pathology. By ‘hierarchical disruption’, we refer specifically to evidence of dysregulation of individual co-expression modules as well as higher-level disruptions in how such modules interact. Our approach is substantially different from previous work in leveraging the identification of consensus modules that robustly exist across datasets. We also account for known methodological differences intrinsic to the existing studies (e.g. age, gender, brain areas). Our approach also utilizes identical parameters to identify consensus networks across datasets, as these parameters vary across different studies in the literature (e.g., network type - signed vs unsigned, soft-power thresholds, deepSplit parameter for cutting dendrograms). These parameters can affect how modules are identified (i.e. clustering and cutting dendrograms) as well as affect the content (i.e. genes, module eigengene variability) composing different discovered modules, thus making direct cross-study comparisons across the literature somewhat difficult. Such a viewpoint of integrating information from multiple datasets currently does not exist in the literature, and we specifically implement this type of analysis in order to be best-positioned to make inferences that are applicable across datasets.

Here, we also directly address ‘replication’ as it pertains to ASD gene expression studies. While some existing studies we have re-analyzed here [20, 21] constitute ‘conceptual’ replications (i.e. studies that differ in numerous ways, yet show roughly similar findings, such as similar gene ontology enrichment results) and are definitely important in their own right, more exact attempts at replication holding constant a variety of analysis issues may also prove insightful. For example, several independent studies may not detect certain dysregulated signals due to analytic or other methodological variance across studies. Repeated detection of such dysregulated signals across multiple studies under more uniform conditions of analysis may likely pull out such prominent signals that otherwise go undetected. Also of importance is how to formally quantify evidence for or against replication. Such formal quantification is missing in the ASD gene expression literature. In this study we tackle these issues head on by analyzing multiple datasets under uniform analysis conditions (e.g., extracting consensus co-expression networks that exist in multiple datasets) and utilize new Bayesian methods developed directly from replication debates ongoing in other fields like psychology, that more formally quantify the strength of evidence for or against replication.

This work also represents the first study to specifically aim at examining hierarchical disruption of the cortical transcriptome in ASD. That is, we go beyond examination of dysregulation at the level of single gene co-expression modules and also examine whether dysregulation is present in higher-level interactions between modules. We provide the first look at the full organization of correlations between gene modules across the transcriptome (i.e. eigengene networks) and examine how such connections manifest differently both at the level of inter-modular connectivity (i.e. connections between specific modules) as well as connectivity relevant to organization of clusters of highly correlated modules (i.e. meta-modules) [22–24]. Localized subtle/specific changes in eigengene network organization or larger global patterns of network reorganization are both plausible predictions regarding how eigengene networks are organized differently in ASD. Both scenarios would lead to the prediction that the composition of meta-modules as well as connectivity within and outside of normative meta-module boundaries would differ in ASD.

## Materials and Methods

### Datasets

We re-analyzed two existing datasets probing cortical gene expression in ASD. The first dataset utilized microarrays on frontal (BA9; n =16 ASD; n = 16 Controls) and temporal cortex (BA 41/42; n = 13 ASD; n= 13 Controls) tissue and was first described by Voineagu and colleagues (Gene Expression Omnibus (GEO) Accession ID: GSE28521) [21]. The second dataset utilized RNAseq on frontal (BA10, n = 6 ASD, n = 8 Controls; BA44, n = 16, n = 11 Controls) and occipital cortex (BA19, n = 24 ASD, n = 38 Controls) tissue and was first described by Gupta and colleagues (http://www.arkinglab.org/resources/) [20]. Across both datasets there was a total n=162 samples (Voineagu n samples = 58; Gupta n samples = 104). The total number of unique individuals across both datasets was n=86 (Voineagu n=32; Gupta n = 54). Of the total n=73 unique individuals within the Gupta dataset, n=19 individuals overlapped with Voineagu dataset. n=26 samples of the total n=104 (25% of samples) from the Gupta dataset came from these overlapping individuals, while the remaining n=76 samples come from new individuals.

For each dataset we utilized the already pre-processed and quality controlled datasets publicly available in order to be as congruent as possible with prior published work. For genes with multiple probes in the Voineagu dataset we selected the probe with the highest mean expression value across the full dataset using the collapseRows function in R [25]. Within the Gupta dataset, missing values were present for some genes in some subjects and these missing values were imputed using the impute.knn function within the impute R library. This procedure was done in order to maximize the total number of genes possible for inclusion into further WGCNA analysis. All further analyses utilize a subset of the 8,075 genes that were common across both datasets.

### Weighted Gene Co-Expression Network Analysis (WGCNA)

Co-expression analysis was implemented with the WGCNA package in R [26]. A consensus WGCNA analysis was implemented in order to detect consensus modules for cross-dataset comparisons (implemented with the blockwiseConsensusModules function) [22]. Consensus WGCNA analysis consisted of construction of correlation matrices, which were then converted into adjacency matrices that retain information about the sign of the correlation (i.e. signed networks use a transformation of 0.5*(r+1)). Adjacency matrices were raised to a soft power threshold selected based on an analysis across various soft power thresholds and choosing the soft power threshold based on a measure of R^2^ scale-free topology model fit that maximized and plateaued well above 0.8 (i.e. soft power = 14 for both datasets; see Fig S1). Soft power thresholded adjacency matrices were then converted into a topological overlap matrix (TOM) and a TOM dissimilarity matrix (i.e. 1-TOM). The TOM dissimilarity matrix was then input into agglomerative hierarchical clustering using the average linkage method. Gene modules were defined from the resulting clustering tree and branches were cut using a hybrid dynamic tree cutting algorithm (the deepSplit parameter was left at the default value of 2) [27]. Modules were merged at a cut height of 0.2 and the minimum module size was set to 30. For each gene module, a summary measure called the module eigengene (ME) was computed as the first principal component of the scaled (standardized) module expression profiles. Genes that cannot be clustered into any specific module are left within the M0 module, and this module is not considered in any further analyses.

To test for differential expression at the level of ME variation we used linear mixed effect models implemented with the lme function in the nlme R library. These models included diagnosis as the fixed effect of interest and additionally included age, sex, RIN, PMI, brain region, and median 5’ to 3’ prime bias (specific to Gupta dataset) as fixed effect covariates. Subject ID was modeled as the within-subject random effect modeled with random intercepts. To identify MEs with replicable differential expression across both datasets, we utilized t-statistics from the linear mixed models to compute replication Bayes Factor (repBF) statistics [28] that quantify evidence for or against replication (see here for R code: http://bit.ly/1GHiPRe). Replication Bayes Factors greater than 10 are generally considered as strong evidence for replication. To identify replicable modules we first considered modules that possessed a significant effect passing FDR [29] q<0.05 within the Voineagu dataset and then also required these modules possess significant effects in the Gupta dataset (FDR q<0.05) and that this evidence quantitatively produces evidence for replication with a replication Bayes Factor statistic > 10.

### Gene-Level Differential Expression Analyses

Differential expression analyses at the level of individual genes were performed in R. The same linear mixed-effect models used for analysis of dysregulation of ME variation were also used for these analysis (e.g., lme function from nlme R library, same fixed and random effects). Genes passing FDR q<0.05 were considered differentially expressed genes. To identify common differentially expressed genes across datasets, we ran gene set overlap analyses implemented using the sum(dhyper()) function in R. The background pool total was set to the total number of genes common to both datasets (8,075).

### Process Level Gene Set Enrichment Analyses

To characterize specific biological processes for all modules, we performed process level (i.e. Process Networks) enrichment analyses within the MetaCore GeneGO software platform. To identify emergent processes from collections of highly correlated dysregulated modules we used GO biological processes enrichment analysis (AmiGO 2; http://amigo.geneontology.org/) in order to leverage GO’s relatively broader hierarchical structure (compared to MetaCore GeneGO). For these enrichment analyses we used a custom background of 7,872 genes which represented all genes analyzed minus the 203 genes within the M0 module. REVIGO [30] was then utilized on the top 50 GO terms ranked by fold enrichment in order to assist in reducing the large number of GO terms into semantically similar clusters of terms. We manually edited the REVIGO output by inserting custom descriptive terms for each cluster and to correct for obvious errors in semantic clustering (e.g., a term like ‘synaptic organization’ occurring outside of the synaptic cluster).

### Cell Type/Cellular Compartment Enrichment Analyses

To characterize differentially expressed modules by enrichments in specific cell types (neuron, astrocyte, oligodendrocyte, M1 and M2 microglia states), and cellular compartments (synapse, postsynaptic density, ribosomal subunits), we utilized lists of markers previously used by Gupta and colleagues [20]. The exception to this was lists of ribosomal subunit markers. These were obtained from lists contained in GO. Enrichment tests were hypergeometric tests (i.e. sum(dhyper()) in R) using the total 7,872 genes as the background pool.

### Eigengene Network Analysis

Eigengene network analysis proceeded by constructing robust ME partial correlation matrices separately for each group. These matrices were computed in MATLAB using robust regression to be insensitive to outliers [31] and the robust regression models incorporated the removal of variation from nuisance covariates (i.e. age, sex, RIN, PMI, median 5’ to 3’ bias, brain region). Partial correlation matrices were then converted into adjacency matrices that retain information about the sign of the correlation. ME adjacency matrices were converted into topological overlap dissimilarity matrices (1-TOM) and then were inserted into agglomerative hierarchical clustering using the ward.D linkage method. The resulting cluster tree was then cleaved into meta-modules using the same dynamic hybrid tree cutting algorithm utilized in WGCNA. We used a deepSplit parameter of 3 since this selection was optimal over and above other options for being able to accurately capture the major branch divisions that are apparent upon visual inspection of the dendrograms.

To visualize eigengene network topology we utilized the qgraph library in R [32] to construct weighted graphs of the ME adjacency matrices for each group. These graphs are depicted using a spring embedded layout algorithm [33] whereby highly connected nodes are attracted to each other and less highly connected nodes are repulsed away from each other. Because these plots are constructed from the adjacency matrices, distance is furthest apart when the correlation is r = -1 and closest when r = 1.

All hypothesis tests on connectivity strength between replicable differentially expressed modules, within and outside meta-module connectivity, and specific intermodular (i.e. between-module) connectivity were implemented with permutation tests (10,000 iterations). The test statistic in each case was the difference in connectivity strength between ASD and Controls. On each iteration we randomized group labels and recomputed the test statistic. FDR [29] q<0.05 was used as the threshold for multiple comparisons correction. Statistically significant results from this analysis are indicated by stars within Figs 7C and 8C and as green outlines around cells within Figs 7D and 8D.

### Protein-Protein Interaction Analysis Between Dysregulated Co-Expression Modules

To further underscore that statistical dependencies in highly correlated coexpression modules indicate direct protein interactions between modules, we implemented a protein-protein interaction analysis. Specifically, if there is hierarchical molecular pathology above single-dysregulated modules indicated by highly interacting dysregulated modules, we should also expect that the degree of such protein interactions would be much higher compared to non-dysregulated and dysregulated module pairings. To test this hypothesis we used Java-based command line tools for GeneMANIA [34] to query the latest protein-protein interaction database (Data Set ID: 2014-08-12; Database Version: 1 June 2014). For each of the 27 modules used as seed modules, we quantified the number of protein-protein interactions between these seed modules and other genes within specific dysregulated module categories (i.e. number of connections between the seed module and downregulated or upregulated modules respectively). If the seed module was itself a dysregulated module, we did not count self-connections (i.e. connections between genes within the same module) in order to guard against results showing higher number of connections simply due to high connectivity within the seed co-expression module. Because co-expression modules differ in size (e.g., the largest module, M1, contains 1568 genes, while the smallest module, M27, contains only 39 genes), we plotted the number of connections for each module as a function of module size. We expect that if dysregulated seed modules are indeed more highly enriched in connections with other dysregulated modules, that the number of connections would be much higher than other non-dysregulated modules of similar size.

## Results

### Replicable Dysregulation of Specific Gene Modules in ASD

Consensus WGCNA on the 8,075 genes common to both the Voineagu and Gupta datasets identified 27 co-expression modules. Information regarding the enrichments for each of these modules can be found in Table S1. Module membership (i.e. the correlation between a gene and its module eigengene) and the top 10 hub genes based on module membership for each module are reported in Table S2. Ten of the 27 modules were identified as differentially expressed in a replicable fashion across datasets (i.e. replication Bayes Factor > 10; see Table S3 for full statistical information on these comparisons). Five of these 10 modules were on-average upregulated in ASD, while the remaining 5 were on-average downregulated in ASD. Three of the 5 ASD-upregulated modules (M12, M24, M27) were enriched in a variety of processes related to the immune system and inflammation; processes such as interferon signaling, complement system, phagocytosis, innate immune response to RNA viral infection, among several others (Fig 1). Interestingly, M12 and M27 are also enriched in M1 microglia markers, while M24 is enriched in M2 microglia markers (Fig 3; Table S4). The ASD-upregulated M25 module was heavily enriched for translation initiation and this enrichment is driven by a large number of genes coding for ribosomal proteins for the 40 and 60S ribosomal subunits (Fig 1). These genes also contributed to a significant enrichment in markers for the postsynaptic density (Fig 3; Table S4). The ASD-upregulated M1 module was also enriched in translation elongation-termination proceeses (Fig 1) and astrocyte and M2 microglia markers (Fig 3; Table S4). In contrast to the ASD-upregulated modules, the replicable ASD-downregulated modules were enriched in a variety of processes that occur at the synapse – GABAergic neurotransmission, synaptic vesicle exocytosis, long-term potentiation, and transmission of nerve impulse (Fig 2). In terms of cell type and cellular component enrichment, downregulated modules are enriched in neuronal (M3, M14), synaptic (M9), and postsynaptic density markers (M9) (Fig 3; Table S4).

**Fig 1:**
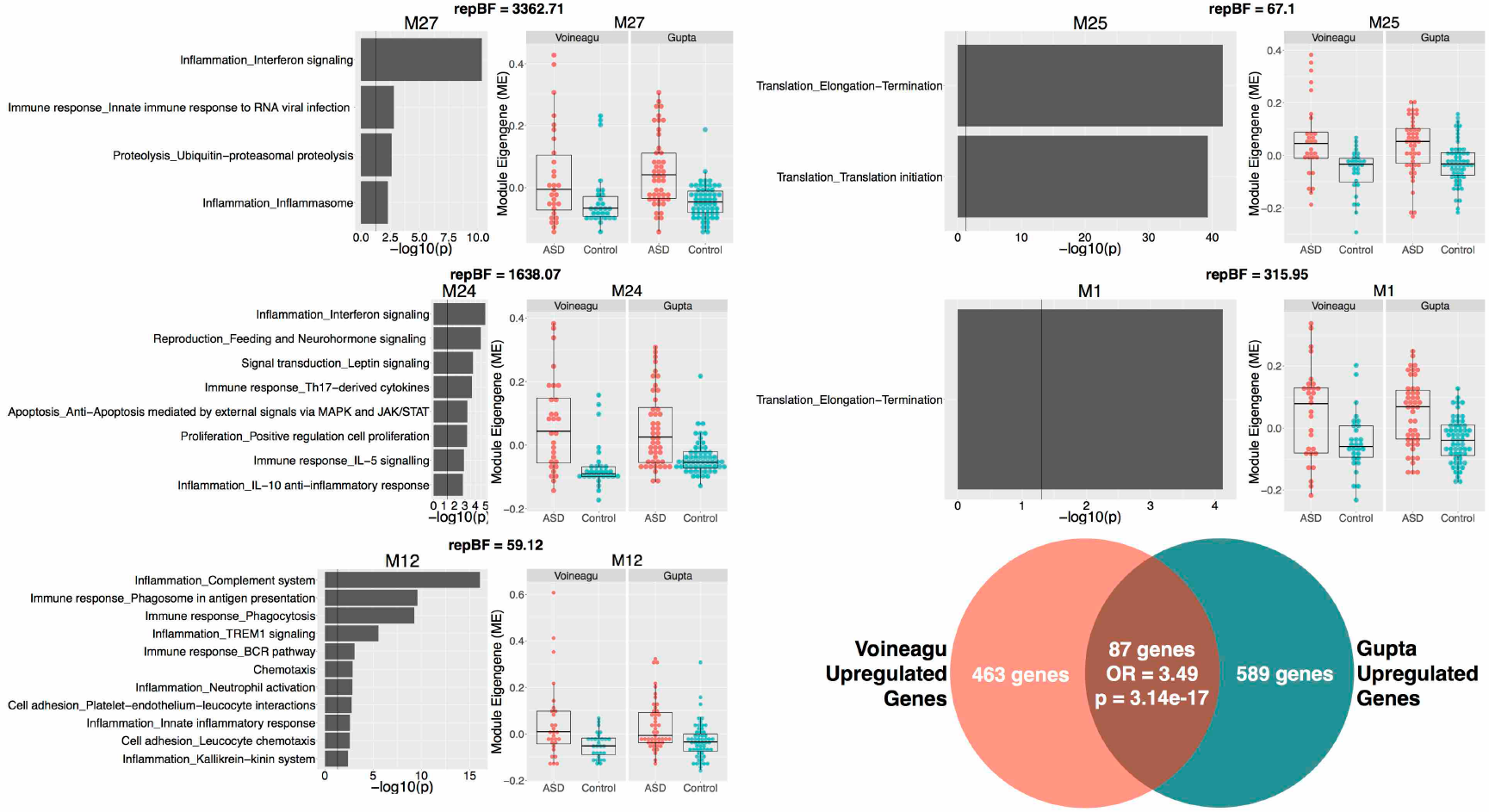
Upregulated gene co-expression modules in ASD.

This figure shows gene coexpression modules that were on-average elevated in ME expression in ASD and in a replicable manner across datasets. Each module has a scatter-boxplot whereby each individual is represented by a dot and the central tendency (median) and dispersion (interquartile range) is shown with the boxplot. Next to each scatter-boxplot are the process-level enrichment terms passing FDR q<0.05 (limited to the top 10 terms) from MetaCore GeneGO. The vertical black line on the enrichment bar plots represents p = 0.05. For each module, the replication Bayes Factor statistic (repBF) is cited above the scatter-boxplot (repBF > 10 indicates strong evidence for replication). In the bottom right corner of this figure is a Venn diagram summarizing the common overlap between ASD-upregulated genes across both datasets.

**Fig 3:**
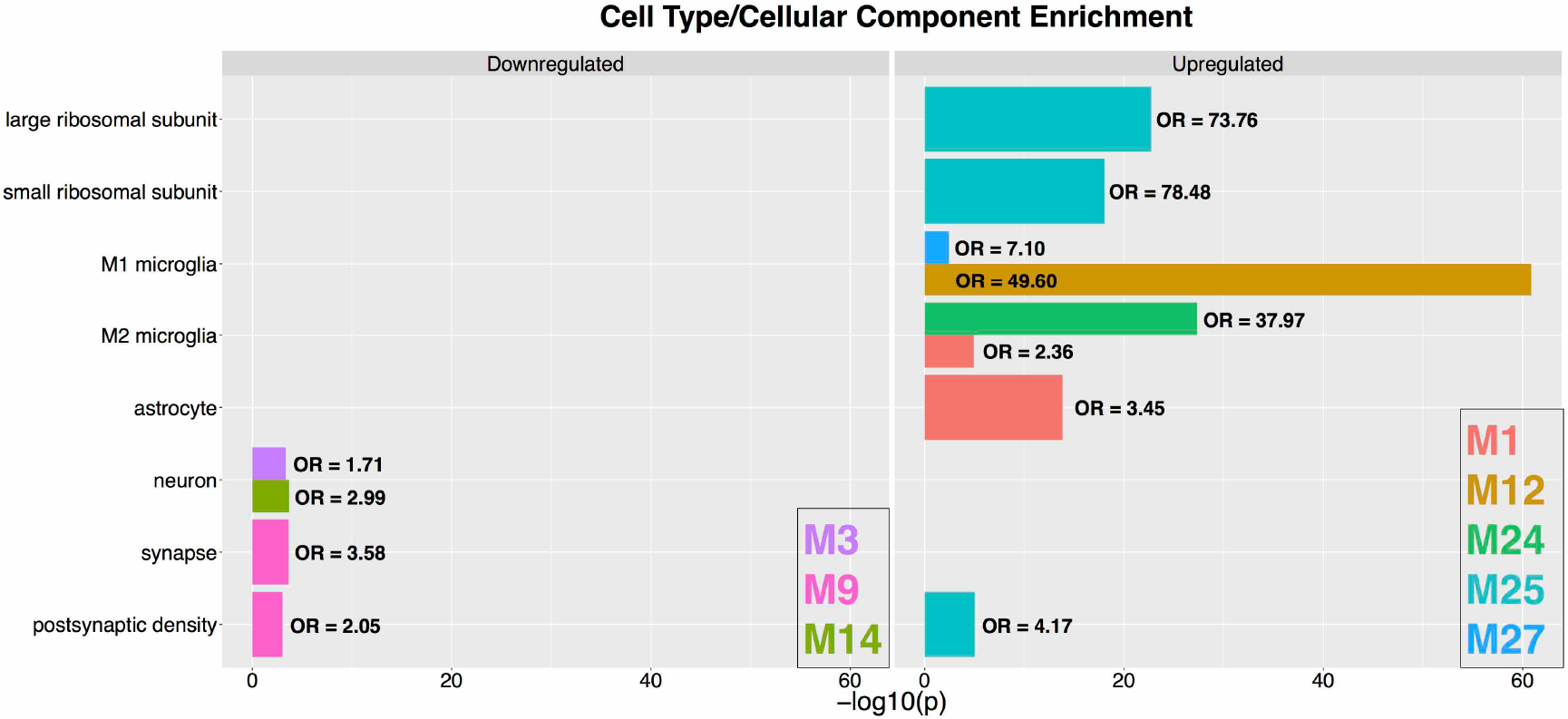
Cell type/cellular compartment enrichments for dysregulated modules.

This figure shows enrichments in a variety of cell types and cellular components for the modules that are replicably dysregulated in ASD. The left panel shows enrichments for downregulated modules, while the right panel shows enrichments for the upregulated modules. The coloring of the bars denote which specific module shows the enrichment and the color legend is shown in the bottom right box for each panel. The x-axis plots the –log10 p-values while the y-axis indicates the specific cell type or cellular compartment. Next to each bar we indicate the enrichment odds ratio (OR).

Results from differential expression analysis at the gene level also showed high degree of overlap between datasets (Fig 1 and Fig 2; Table S3). We utilized these gene level differential expression results to characterize the load of differential expression signal within each of the 10 identified replicable dysregulated modules. Congruent with the labels of ‘upregulated’ or ‘downregulated’ modules, we find that each of these modules are heavily loaded with differential expression signal at the gene level congruent with such labels (Fig S2). For example, all ‘upregulated’ modules had pronounced shifts in differential expression signal in the direction of ASD>Control, while all ‘downregulated’ modules had pronounced differential expression signal shift in the direction of Control>ASD. These results confirm that the on-average between-group differentiation in dysregulated modules is underpinned by large magnitude of differential expression signal within each module at the level of individual genes.

**Fig 2:**
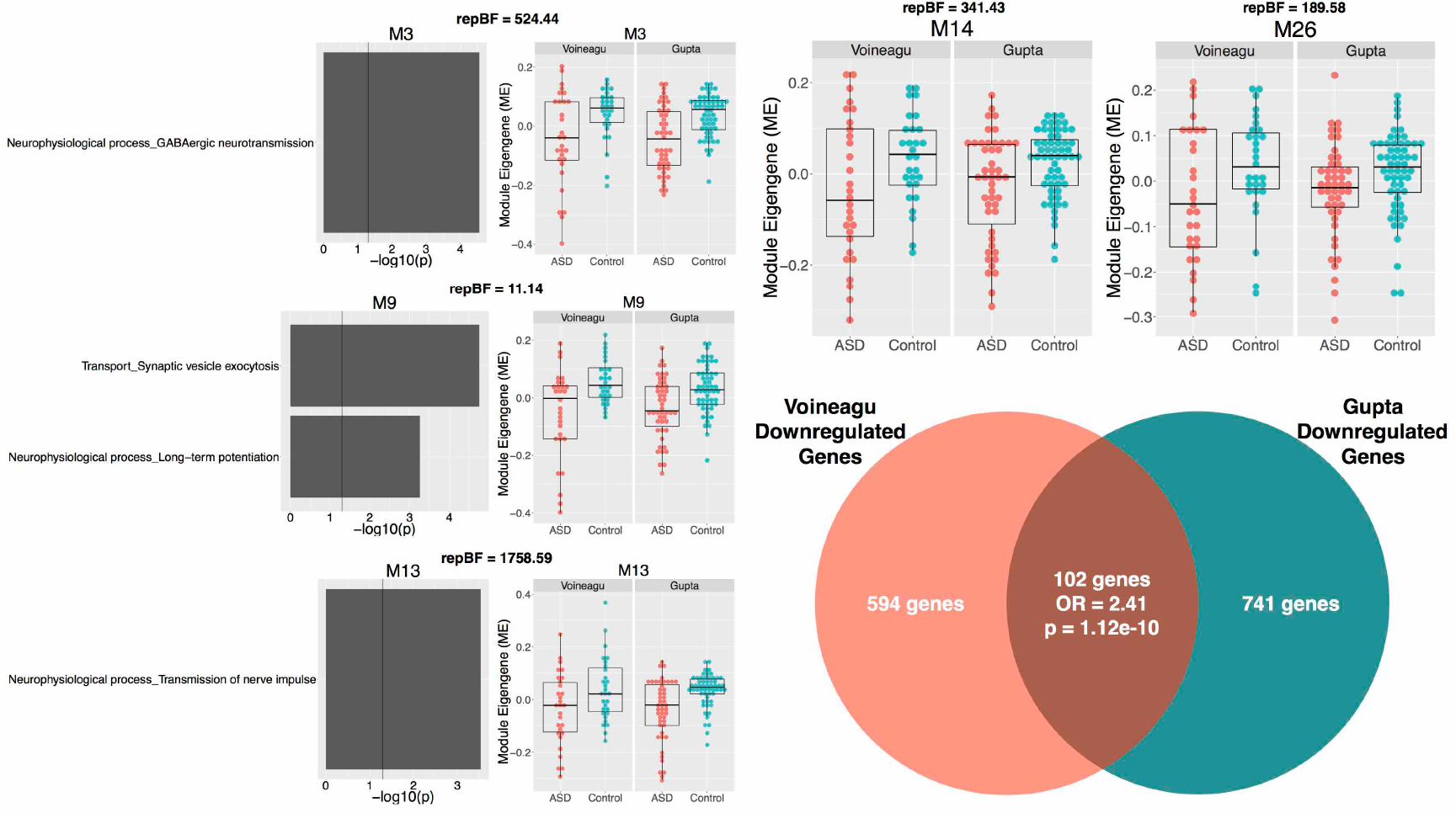
Downregulated gene co-expression modules in ASD.

This figure shows gene co-expression modules that were on-average decreased in ME expression in ASD and in a replicable manner across datasets. Each module has a scatter-boxplot whereby each individual is represented by a dot and the central tendency (median) and dispersion (interquartile range) is shown with the boxplot. Next to each scatter-boxplot are the process-level enrichment terms passing FDR q<0.05 (limited to the top 10 terms) from MetaCore GeneGO. The exception here is M26, whereby none of the terms passed FDR q<0.05. In this instance, we plot the first 5 terms for descriptive purposes. The vertical black line on the enrichment bar plots represents p = 0.05. For each module, the replication Bayes Factor statistic (repBF) is cited above the scatter-boxplot (repBF > 10 indicates strong evidence for replication). In the bottom right corner of this figure is a Venn diagram summarizing the common overlap between ASD-downregulated genes across both datasets.

### Differentially Expressed Modules are Highly Correlated in ASD

Modules that are on-average differentially expressed (Figs 1–2) are highly correlated. This pattern of correlation was one of strong positive correlations within modules that share similar directionality of differential expression, but strong negative correlations between modules with different directionality of differential expression. Interestingly, these correlations become significantly enhanced in ASD compared to Controls in the Voineagu dataset (within downregulated modules p = 0.012; within upregulated modules p = 0.042; between downregulated and upregulated modules p = 0.008; Fig 4A-B). Within the Gupta dataset, this phenomenon of highly correlated differentially expressed modules as well as strong negative correlations between upregulated and downregulated modules is already present in Controls and stays present in ASD, though quantitative strengthening of such connectivity in ASD does not occur (within downregulated modules p = 0.957; within upregulated modules p = 0.327; between downregulated and upregulated modules p = 0.667; Fig 4C-D).

**Fig 4:**
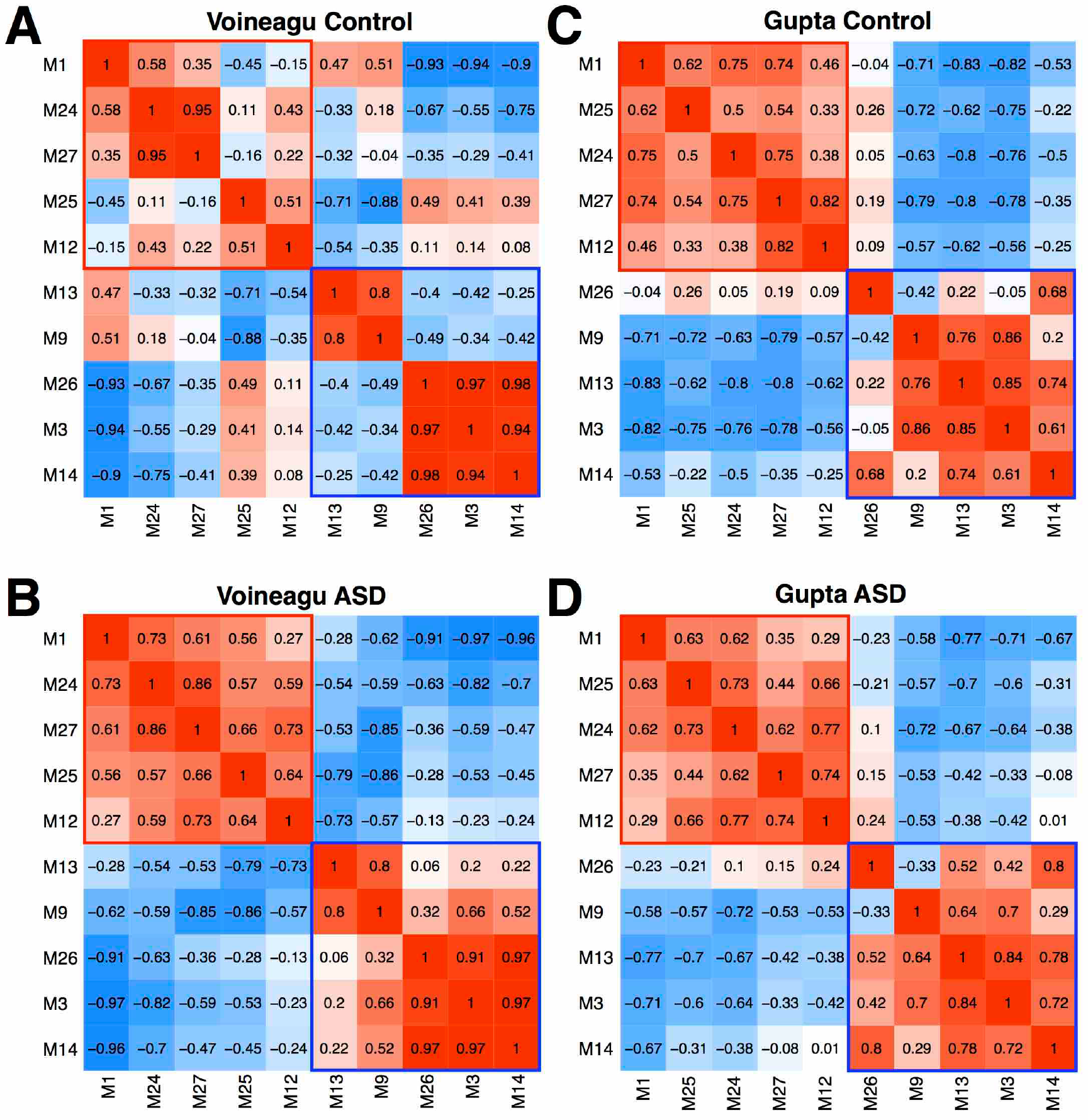
Correlations between dysregulated modules.

Panels A and B show correlations between differentially expressed modules in the Voineagu Control (A) or ASD (B) datasets. Panels C and D show correlations between these same modules in the Gupta Control (A) or ASD (B) datasets.

### Highly Correlated Differentially Expressed Modules Highly Interact at the Level of Protein-Protein Interactions

Statistical dependencies (i.e. correlations) between dysregulated co-expression modules suggest that hierarchical pathology may be present in the interactions between such modules. From this result, we further reasoned that strong correlations between dysregulated modules may result from high levels of direct physical interactions between proteins of such modules. If some important synergistic pathology were apparent across such ASD-dysregulated modules we would also expect that the high degree of protein-protein interactions between collections of dysregulated modules would be much stronger degree of interactions between dysregulated and non-dysregulated modules. Such protein-protein interaction evidence would further indicate plausibility of the idea that hierarchical pathology evident in the interactions between dysregulated co-expression modules exists in ASD. To answer this question, we queried the GeneMANIA protein-protein interaction database [34] and discovered that each of the dysregulated modules do indeed show a large degree of connections with other dysregulated modules. Interestingly, modules dysregulated in opposite directions (e.g., connections between a downregulated seed module and all other upregulated modules) showed just as many connections as modules dysregulated in the same direction. While one might expect high degree of connections between modules dysregulated in the same direction (e.g., modules dysregulated and enriched in similar kinds of biological processes), the fact that similar numbers of protein interactions exist between modules dysregulated in opposite directions (e.g., connections between a downregulated synaptic seed module and other upregulated immune and translation modules) supports the idea that large-scale hierarchical interactions are important to the pathophysiology of ASD. Importantly, the number of connections between dysregulated seed modules and other dysregulated target modules was much higher than when non-dysregulated modules were the seed. This is evident in the predicted observation that dysregulated seed modules show much higher degree of connections than non-dysregulated seed modules of similar size (Fig 5). This evidence alongside the observed statistical dependencies between co-expression modules further support the idea that disparate co-expression modules enriched in different biological processes are likely interacting in important ways in the pathophysiology of the ASD cortical transcriptome.

**Fig 5:**
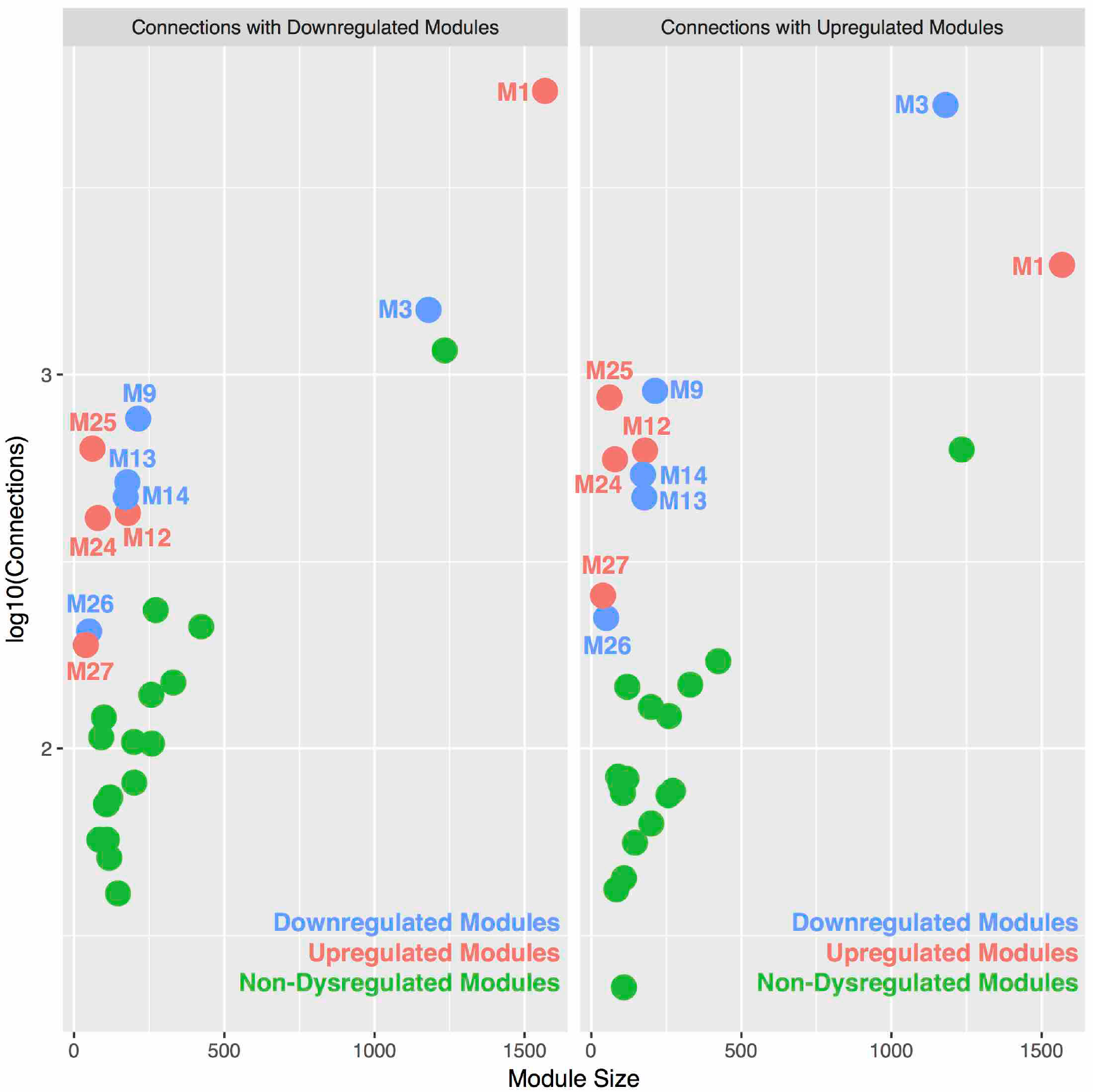
Protein-protein interactions between dysregulated modules.

This figure plots the number of protein-protein interactions (on log10 scale) between seed modules and downregulated (left) or upregulated (right) modules (y-axis) as a function of module size (number of genes in the module; x-axis). Seed modules that are downregulated are colored in blue, while upregulated seed modules are colored in red. Non-dysregulated seed modules are colored in green. For dysregulated seed modules, the number of connections reflects the number of protein connections with other dysregulated modules, not counting self-connections (e.g., connections between genes of the same co-expression module). This figure clearly shows that seed modules that are dysregulated (red or blue) possess a far greater number of connections with other dysregulated modules compared to non-dysregulated modules (green) of a similar size.

### Processes Enriched within Dysregulated Modules

We next asked the question of what biological processes might characterize such emergent phenomena of interacting collections of co-expression modules. Leveraging the hierarchical structure of Gene Ontology (GO), we input merged lists of all differentially expressed modules together and computed GO biological process gene set enrichment and then clustered the top 50 enriched GO terms by semantic similarity [30]. Here we find that the emergent process represented by the combination of highly connected downregulated modules is primarily involved in synaptic function (Fig 6A). In contrast, there were several emergent processes represented by the combination of highly connected upregulated modules – immune/inflammation processes, response to other organism, viral processes, catabolism, translation, protein targeting, and localization, cell proliferation, and vasculature development (Fig 6B). These results suggest that highly connected differentially expressed modules spanning multiple cell types and cellular compartments, also interact at the protein level and result in emergent phenomena that are not visible simply by examining modules in isolation.

**Fig 6:**
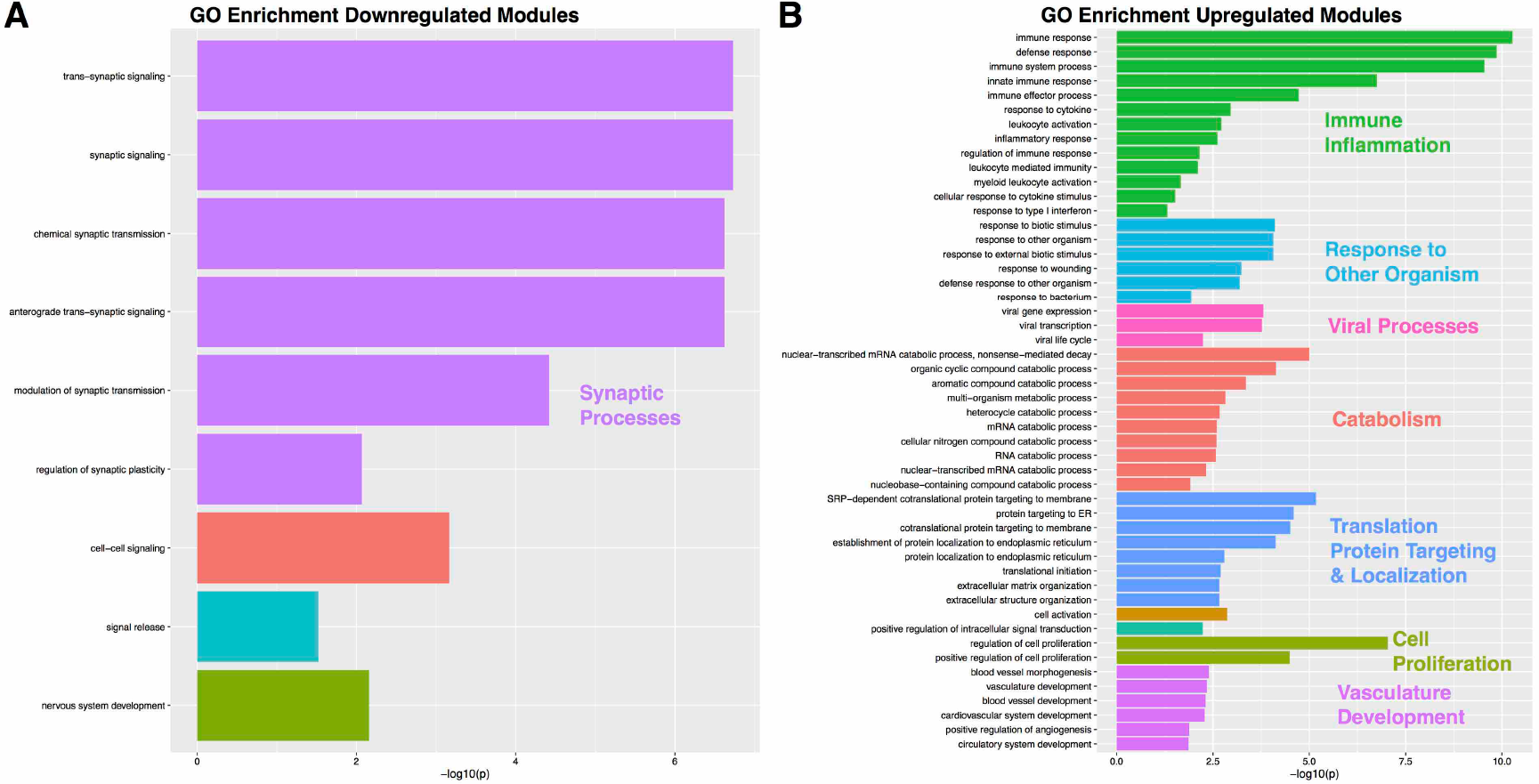
GO biological process enrichments for collections of downregulated or upregulated modules.

This plot shows GO biological process enrichment terms for the combination of all downregulated (A) or upregulated (B) modules. The top 50 GO terms ranked by fold enrichment were input into REVIGO [30] in order to cluster GO terms by semantic similarity. These clusters are shown in different colors along with a descriptive label for each cluster. Plotted on the x-axis of each plot is the Bonferronicorrected –log10 p-value for each term.

### Topological Reorganization of Eigengene Networks

While we have primarily focused on dysregulated modules, viewing hierarchical organization just within these 10 modules limits the insights that could be made by examining the full hierarchical organization of eigengene networks across all modules. By examining eigengene networks, we can observe organization at higher levels above individual co-expression modules. Such observations can show how individual modules cluster into collections of highly connected modules, known as ‘meta-modules’. These analyses go beyond the 10 individually dysregulated modules to allow for further insights into how eigengene networks are composed across groups and to provide deeper insights into how such organization takes shape with regards to meta-modular clustering. These analyses visualize full eigengene network organization and meta-module membership with spring-embedded graphs that indicate topological change via distancing nodes based on strength of correlation between modules (i.e. shorter distance indicates stronger correlation, further distance indicates weaker correlation). We also quantitatively tested for differences with respect to connectivity strength within and outside meta-module boundaries as well identifying specific modules with disrupted connectivity.

Initial examination of preservation of the Control eigengene network organization indicated that high levels of preservation are not present across many nodes of the network (see Fig S3). Thus, this low level of preservation suggests that assessing replicability of any between-group differences in eigengene network organization across datasets is likely not possible. Unlike our initial consensus WGCNA analysis that ensured similar co-expression networks across datasets, this procedure does not guarantee that eigengene network organization may be similar, and this analysis verifies that the organization of eigengene networks across datasets differs considerably. This effect could be due to a variety of the methodological factors that differ across these datasets (e.g., different brain regions, differing age of the samples, microarray vs. RNA-seq, etc). Nevertheless, this issue does not invalidate observations of how eigengene network organization differs within each dataset, and therefore, we restrict our descriptions of eigengene network organization to each dataset independently.

Within the Voineagu dataset, ASD-dysregulated modules are topologically arranged closer together in ASD and within the same meta-module, compared to the more disperse and heterogeneous organization in Controls with respect to metamodule membership of dysregulated modules. This differing pattern of topological organization at the meta-module level can be clearly seen in the spring-embedded graph layouts shown in Fig 7A-B. For example, upregulated modules (circled in red in Fig 7A-B) are spread across 3 different meta-modules in Controls, while in ASD these modules are positioned close together and within the same orange meta-module (Fig 7A-B). Quantitatively, network reorganization can be examined in connectivity strength differences within and outside normative (Control-defined) meta-module boundaries. Four modules (M25, M9, M21, and M23) show ASD-decreased connectivity within normative meta-module boundaries. These same modules along with one other module (M16) also show enhanced connectivity outside of normative meta-module boundaries in ASD (Fig 7C). At a nodal level, we further observed specific between-module connections that are prominently affected in ASD (Fig 7D). The ASD-upregulated M25 translation initiation module is normatively negatively correlated with the prominent ASD-upregulated M27 interferon signaling and M1 translation elongation-termination module. However, in ASD, these negative correlations significantly reverse and turn into positive correlations, suggesting some abnormally heightened integration between these distinct biological processes/pathways. In another example, the ASD-downregulated M9 module is normatively positively correlated with M1, M15, and M16, but these relationships reverse into negative correlations in ASD. This suggests that what should typically be a natural integration between these modules ends up being an abnormal lack of integration in ASD. Furthermore, M9’s connectivity with another ASD-downregulated module (M3) is normatively negative, yet in ASD is highly positively correlated. Finally, while there is little to no normative relationship between the ASD-downregulated M9 module and the ASD-upregulated M27 module, in ASD this relationship turns into a strong negative correlation. This effect could potentially indicate an abnormal immune-synapse interaction between upregulation of inflammation interferon signaling processes and downregulation of important synaptic processes in ASD.

**Fig 7:**
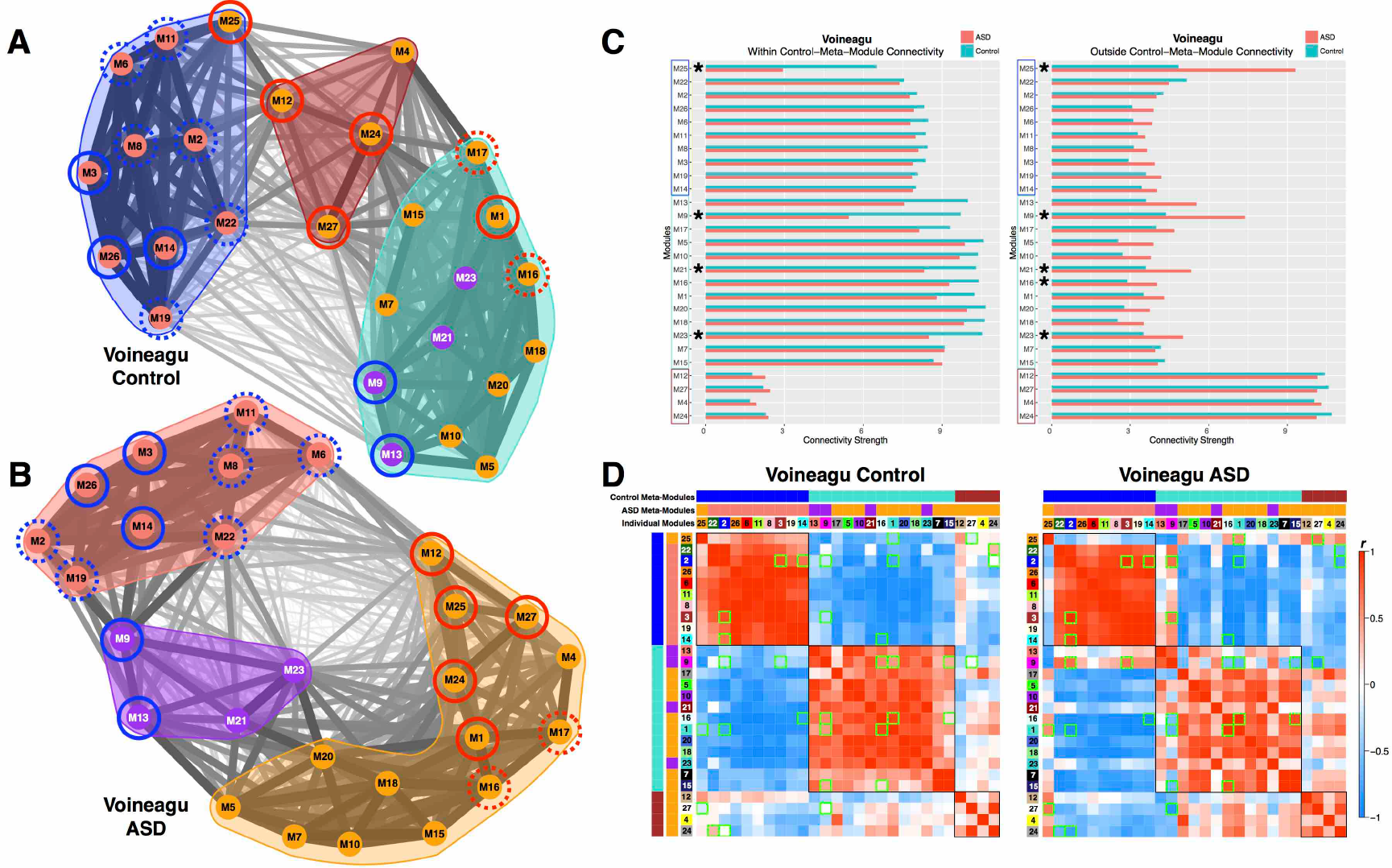
Eigengene network topology and connectivity differences within the Voineagu dataset.

Panels A and B show eigengene networks as weighted graphs in a spring embedded layout for the Voineagu Control (A) or ASD (B) groups. The spring embedded layout places modules (nodes within the graphs) that are highly connected as much closer in space whereas modules that are less highly connected are repelled away from each other. The thickness of the connections (i.e. edges) between modules are scaled to connection strength whereby the thinnest line represents a correlation of r = -1 and the thickest line represents a correlation of r = 1. The color of each module node represents the ASD meta-module it belongs to. This was done to represent where the ASD meta-modules are located within the Control graph. The color-filled outlines around collections of modules represent the meta-module boundaries. Modules with a solid red or blue circle around it are modules that were identified in Figs 1–2 as being replicably dysregulated in ASD across both datasets (blue = ASD-downregulated; red = ASD-upregulated). The dotted circles represent differentially expressed modules (FDR q<0.05) present only within that specific dataset (see Table S3). Panel C shows within (C) and outside (D) normative metamodule connectivity strength for each seed module depicted on the y-axis. The normative (Control-defined) meta-modules are denoted by the color of the rectangular outlines on the y-axis. Connectivity strength is depicted on the x-axis and for within meta-module connectivity is defined as the sum of connection strength between the seed module and all other modules within the seed module’s normative meta-module. Outside meta-module connectivity strength is defined as the sum of connection strength between the seed module and all other modules outside of the seed module’s normative meta-module. Turquoise bars indicated Controls and salmon colored bars indicate ASD. The stars next to specific modules indicate a significant between-group difference in connectivity strength. Panel D illustrates eigengene networks as robust ME partial correlation matrices. Red coloring within the matrices indicates increasing positive correlation strength, while blue coloring indicates increasing negative correlation strength; see colorbar for key indicating how color corresponds to correlation strength. Matrices have rows and columns ordered by hierarchical clustering based on the Control group and the individual module numbers as well as meta-module colors are shown. Normative (Control-defined) meta-module boundaries are also clearly delineated by the black outlines over cells in the correlation matrices. Any cells with green outlines are those specific between-module connectivity comparisons that differed between-groups.

Within the Gupta dataset there was also evidence of topological reorganization, with a much more fractionated organization of meta-modules in ASD compared to Controls (i.e. 6 meta-modules in ASD versus 4 in Controls). This differing pattern of topological organization at the meta-module level can be clearly seen in the spring-embedded graph layouts shown in Fig 8A-B. Similar to the Voineagu dataset, dysregulated modules again clustered close together and within the same meta-modules relative to a more heterogeneous organization in Controls (Fig 8A-B). Quantitatively, connectivity within and outside of normative meta-module boundaries was perturbed in ASD for nearly every single module (Fig 8C). This indicates that ASD eigengene network organization is highly perturbed with regard to connectivity of modules within normative eigengene network topology. In contrast to the numerous modules showing connectivity differences at the nodal level in the Voineagu dataset, very few nodal-level differences emerged in the Gupta dataset. Thus within the Gupta dataset, it appears that overall eigengene network topology is reorganized in ASD in subtle ways that are spread across many modules and considerably affect meta-modular organizational structure. However, they cannot be tied to very pronounced and specific differences within specific subsets of modules.

**Fig 8:**
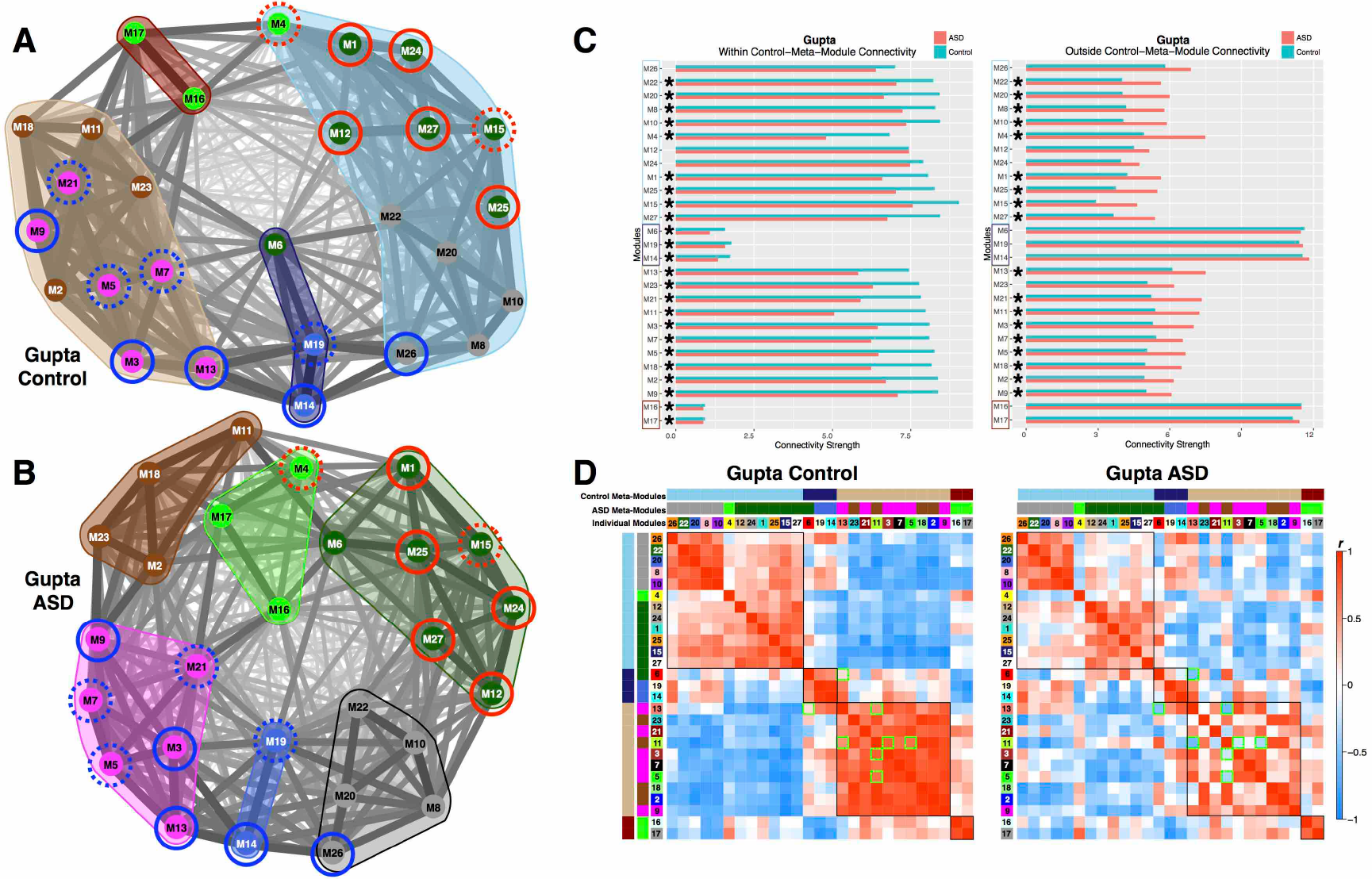
Eigengene network topology and connectivity differences within the Gupta dataset.

Panels A and B show eigengene networks as weighted graphs in a spring embedded layout for the Gupta Control (A) or ASD (B) groups. The spring embedded layout places modules (nodes within the graphs) that are highly connected as much closer in space whereas modules that are less highly connected are repelled away from each other. The thickness of the connections (i.e. edges) between modules are scaled to connection strength whereby the thinnest line represents a correlation of r = -1 and the thickest line represents a correlation of r = 1. The color of each module node represents the ASD meta-module it belongs to. This was done to represent where the ASD meta-modules are located within the Control graph. The color-filled outlines around collections of modules represent the meta-module boundaries. Modules with a solid red or blue circle around it are modules that were identified in Figs 1–2 as being replicably dysregulated in ASD across both datasets (blue = ASD-downregulated; red = ASD-upregulated). The dotted circles represent differentially expressed modules (FDR q<0.05) present only within that specific dataset (see Table S3). Panel C shows within (C) and outside (D) normative meta-module connectivity strength for each seed module depicted on the y-axis. The normative (Control-defined) meta-modules are denoted by the color of the rectangular outlines on the y-axis. Connectivity strength is depicted on the x-axis and for within meta-module connectivity is defined as the sum of connection strength between the seed module and all other modules within the seed module’s normative meta-module. Outside meta-module connectivity strength is defined as the sum of connection strength between the seed module and all other modules outside of the seed module’s normative meta-module. Turquoise bars indicated Controls and salmon colored bars indicate ASD. The stars next to specific modules indicate a significant between-group difference in connectivity strength. Panel D illustrates eigengene networks as robust ME partial correlation matrices. Red coloring within the matrices indicates increasing positive correlation strength, while blue coloring indicates increasing negative correlation strength; see colorbar for key indicating how color corresponds to correlation strength. Matrices have rows and columns ordered by hierarchical clustering based on the Control group and the individual module numbers as well as meta-module colors are shown. Normative (Control-defined) meta-module boundaries are also clearly delineated by the black outlines over cells in the correlation matrices. Any cells with green outlines are those specific between-module connectivity comparisons that differed between-groups.

## Discussion

Here we provide the first detailed characterization of how the ASD cortical transcriptome is hierarchically disorganized both at the level of specific co-expression modules and at higher levels of eigengene network organization (i.e. connectivity between modules and meta-modules). We have pinpointed several novel coexpression signals that show strong evidence for replicable dysregulation across datasets [20, 21]. Rather than pinpointing a single synaptic or immune-related module, we have identified several dysregulated synaptic and immune modules. These modules are differentiated in terms of cell type/compartment enrichment and/or show different biological process enrichment within the broader class of synaptic and immune-related processes. For example, while both M3 and M9 modules are downregulated in ASD and enriched in similar synaptic processes, their cell type/compartment enrichments differ. M3 is primarily enriched in neuronal markers, whereas M9 is specifically enriched in synaptic and postsynaptic density markers. Synaptic M3 and M9 modules also differentiate in how they interact with other modules (see Fig 7C-D for example). These results provide an example of how subtle distinctions may be present within the class of downregulated synaptic signals.

We have also identified multiple types of ASD-upregulated immune/inflammation modules that are novel distinctions from past work. Although prior work has implicated interferon signaling, particularly with respect to M2 microglia markers [20], here we find evidence for 2 upregulated interferon signaling modules (M24, M27). These modules differentiate by M1 and M2 microglia activation states, with M27 enriched in M1 microglia markers while M24 is enriched in M2 microglia markers. Between-module connectivity evidence also suggests that these two interferon signaling modules are disrupted in different ways. M27 is abnormally connected to an important ASD-upregulated translation initiation (M25) and ASD-downregulated synaptic module (M9). Given the enrichment in M27 for M1 microglia activation markers, this evidence suggests that cytotoxic M1 microglia processes may be affecting synaptic proteins in ASD. On the other hand, M24 shows intact connectivity between M25 and M9, but aberrant connectivity between other modules (M2, M22). These results suggest that while upregulated interferon signaling can be linked to both M1 and M2 microglia phenotypes, such aberrant processes may have differing impact on ASD brain function and structure.

In addition to the multiple dysregulated interferon signaling modules, we have also uncovered novel evidence for ASD-upregulation of an immune/inflammation module (M12) enriched in the complement system and phagocytosis processes and M1 microglia markers. In conjunction with effects from interferon signaling modules, the addition of the complement system may be of particular importance given the known links between the complement system and synaptic pruning [35, 36] and remodeling as well as enhancing pro-inflammatory states of microglia activation in ASD [37–39]. Recently, the complement system has been noted as a prominent player in the pathophysiology of schizophrenia, particularly for its role in synaptic pruning [40]. In the larger context of eigengene networks it is interesting that all of these important immune/inflammation modules are members of the same meta-module in ASD and that such a meta-module also includes other prominent modules such as the ASD-upregulated M25 translation initiation module. The current data present a role for complement system signaling alongside interferon signaling and other immune processes working together and potentially in concert with other important modules relating to translation and also for their role in various types of microglia activation states.

New modules not highlighted at all by prior work were also identified. Two of these modules (M1 and M25) are heavily enriched in translation initiation and translation elongation-termination processes and are enriched in genes coding for proteins that make up the 40S and 60S ribosomal subunits (*RPL* and *RPS* genes). Translation has been an important topic in ASD primarily because of work on syndromic forms of autism related to mutations in *FMR1*, *TSC1/2*, and *PTEN* [6, 41], as well as the important cap-dependent translation gene *EIF4E* [42–45]. However, none of this work has specifically implicated ribosomal proteins themselves and no prior work on the cortical transcriptome in ASD has specifically implicated upregulation of translation initiation signals. These modules were dysregulated with respect to connectivity within and outside of normative meta-modular boundaries and showed specific abnormal interactions with each other as well as other ASD-upregulated modules (e.g., M27). Additionally, these translation modules were also a member of a meta-module in ASD that was composed of other upregulated immune/inflammation modules (M12, M24, M27), suggesting that they may play important roles integrating with upregulated immune/inflammation processes in ASD. Thus, not only have we discovered evidence for a novel and important upregulated signal in the ASD cortical transcriptome, but this finding also may have important implications with regards to its potential as a cross-cutting influence on other pathophysiological processes in ASD.

This novel finding of upregulated translation initiation and elongation-termination processes in ASD is important, as it agrees with other work on blood transcriptome markers. Our recent work on blood leukocyte gene expression has also uncovered upregulated translation initiation as a prominent signal in young toddlers with ASD and this signal is present alongside other upregulated immune/inflammation signals, particularly interferon signaling and phagocytosis [46]. Further bolstering these inferences, a recent mega-analysis of seven different studies in the literature and also found ribosomal translation as one prominent upregulated process in blood [47]. The presence of these dysregulated and highly connected translation initiation and immune/inflammation signals across brain and blood is potentially important because it may signal a unique opportunity to assay brain-relevant dysregulation in peripheral tissues and *in-vivo* in living patients. This peripheral window into potentially brain-relevant dysfunction that can be assayed in living patients may be particularly important given the recent discovery of a direct linkage between the brain and lymphatic vessels of the immune system [48]. Investigating this possible peripheral linkage to brain-relevant dysfunction in living patients using *in-vivo* techniques like functional and structural neuroimaging [49] will be an important next step in understanding whether peripherally dysregulated signals in blood play some role in linking directly to important macro-level neural systems dysfunction in living patients [50]. We have also recently identified similar upregulation of translation initiation signals, particularly ribosomal proteins, in a rodent model of maternal immune activation [51], indicating that sources of translation initiation upregulation in ASD may have pathophysiological impact in early fetal development and can be influenced by environmental factors. Furthermore, modeling the upregulated expression of a long non-coding RNA, *MSNP1AS*, in neural progenitor cells also leads to differential expression of genes involved in translation, protein synthesis and which are localized to the ribosome [52]. *MSNP1AS* was the first GWAS hit in ASD [53] and is known to be upregulated in expression in ASD cortex [54]. Influence via this common variant may further indicate influence over this process of translation in the developing ASD brain. Future work using *in-vivo* and *in-vitro* models targeting these novel ribosomal protein genes from the M25 translation initiation module (e.g., hub genes shown in Table S2) may be important for leading to further insights on the pathophysiology behind ASD.

In addition to implicating several new gene co-expression modules of significance to ASD, this work provides primary evidence supporting the idea that the cortical transcriptome is dysregulated at hierarchical levels and this hierarchical view of pathophysiology cannot be well understood from the vantage point of examining single co-expression modules in isolation. By identifying disruption in the interaction between-modules and in how eigengene networks are reconfigured into different meta-modular structures, this work presents a larger view on how multiple dysregulated signals may operate in conjunction with one another and potentially implicate important emergent interactions at the protein level. We show that a number of specific modules that are on-average up- or downregulated in ASD are also highly correlated and that this correlation can become stronger in ASD. This result is not apparent in prior work on this topic, with the closest result being the previous observation of a negative correlation when collapsing across both groups between single pair of modules enriched in synaptic and immune functions [8, 20]. We have gone much further to show correlations between dysregulated modules including translation initiation modules and several other modules. We also demonstrated that beyond the statistical dependencies between co-expression modules, these dysregulated modules physically interact at the level of proteins. The disruption of these coordinated higher-order interactions at a protein level suggests that systems-level phenomena are disrupted in ASD that coordinates disparate biological processes and which cannot be adequately characterized by viewing smaller elements (e.g., single genes, single co-expression modules) in isolation. Thus, a primary conceptual advance from this aspect our work suggests that we may need to move beyond arguments about single unitary processes, since the interactions between multiple dysregulated processes may underlie and better describe the pathology.

As a whole, the collection of ASD-downregulated modules appears to involve a number of processes occurring at the synapse. While synaptic processes are commonly discussed as important mechanisms [21, 55], genes that are typically characterized as synaptic genes may have other pleiotropic roles in very early neural developmental processes. It is known that annotations in enrichment databases (e.g., GO, MetaCore) may be incomplete and an example of this can be seen in potential other interpretations of genes typically thought of as involved in synaptic processes. Casanova and colleagues recently showed that many high-risk ASD genes that have canonical roles in synapse development are also involved in very early stages of neural proliferation, growth, and maturation [56]. As a specific example of this idea, Konopka and colleagues discovered that *NRXN3* plays a role in earlier neural progenitor biology that is different from its later function at the synapse [57]. Early fetal brain developmental processes occurring as early as the end of the first trimester of gestation [58] and are key signals of importance highlighted by prior studies on very early pathophysiology in ASD [5, 12, 17, 49, 56, 59–62]. These specific neural developmental processes are developmentally prior to abnormalities in synaptic processes which emerge at later points in fetal development and continue to change throughout life as a result of postnatal experience and adaptation. Therefore, a nuanced interpretation of the role of synapse gene dysregulation in ASD could be that these genes have pleiotropic roles in both early stages of neural development (e.g., proliferation, growth, and maturation) and at later stages dealing with synaptic processes that continue throughout the lifespan.

The collections of modules upregulated in ASD showed evidence for several novel and emergent biological phenomena. To our knowledge, the novel signal of upregulated catabolism has not been implicated in any past work. Additionally, there are novel upregulated processes involved in protein targeting and localization that can be intertwined with translation processes (e.g., SRP-dependent co-translational protein targeting to membrane). Finally, we also found enrichment in several viral processes, responses to other organisms, cell proliferation, and vasculature developmental processes are highly prominent. These highly coordinated processes are associated with multiple cell types/compartments, and the downregulation of synaptic processes – as evidenced by the strong negative correlations between upregulated and downregulated modules. This evidence is generally in agreement with past theoretical ideas [60] that suggested that early manifestations of pathophysiology potentially emerging in fetal development could then trigger a later corrective phase of development characterized by downregulation of synaptic and neuronal processes and potential upregulation immune/inflammation (e.g., microglia activation) [37–39], apoptotic, and other processes. A challenge for future research will be to unpack the relationships between known and novel upregulated processes with downregulated synaptic and neural developmental processes. However, it is important to underscore that these inferences emerge from the looking at the highly-coordinated interactions between multiple dysregulated co-expression modules, and are not obvious by simply targeting specific modules and looking at such elements in isolation. Thus, these new insights regarding systems level phenomena in ASD can further guide future studies to unravel specific novel mechanisms (e.g., targeting hub genes for many of the dysregulated modules we have implicated and examining their impact on other connected systems-level processes; Table S2).

## Conclusions

In summary, this work highlights several novel aspects about how the cortical transcriptome is dysregulated in ASD. A primary advance of this work is the idea that dysregulation of the cortical transcriptome in ASD does not occur only at the level of individual gene co-expression modules. Rather, the cortical transcriptome is disorganized at higher levels of analysis such as the interactions between modules and how modules form hierarchical organization structure as meta-modules within eigengene networks. The insight that this new view may shed on the biology of autism is yet to be explored, but at the very least implicates that emergent pathology may arise out of interactions across otherwise disparate separate biological processes and pathways. As development progresses, the brain in ASD may be engaging in adaptive processes to compensate for inherent biological problems that originate in very early fetal or postnatal brain development [60, 63, 64]. This might lead to the interesting proposition that the core symptomatology of ASD present by 2-4 years of age, is the direct output of this postnatal early developmental adaptation process that attempts to compensate for early fetal abnormalities in how the brain lays down core elements to build upon with further experience. Our approach here may provide a better viewpoint on how to describe such processes and may further help enable future translational insights.

## Ethics approval and consent to participate

Not applicable

## Consent for publication

Not applicable

## Availability of data and materials

The data utilized in this work was downloaded from publicly available sources. Specifically, the Voineagu et al., data can be found on Gene Expression Omnibus (GEO; https://www.ncbi.nlm.nih.gov/geo/; Accession ID: GSE28521). The Gupta et al., data be found at http://www.arkinglab.org/resources/.

## Author Contributions

MVL, EC, and TP conceived the idea for the study. MVL, NEL, and TP conceived all analyses. MVL implemented all data analyses. MVL, EC, NEL, and TP interpreted the results and wrote the manuscript. All authors read and approved the final manuscript.

## Acknowledgments

We would like to thank Bhismadev Chakrabarti, Bonnie Auyeung, Meng-Chuan Lai, Simon Baron-Cohen, and Peter Langfelder for helpful discussion on this work.

## Supplementary Figures

**Fig S1:**
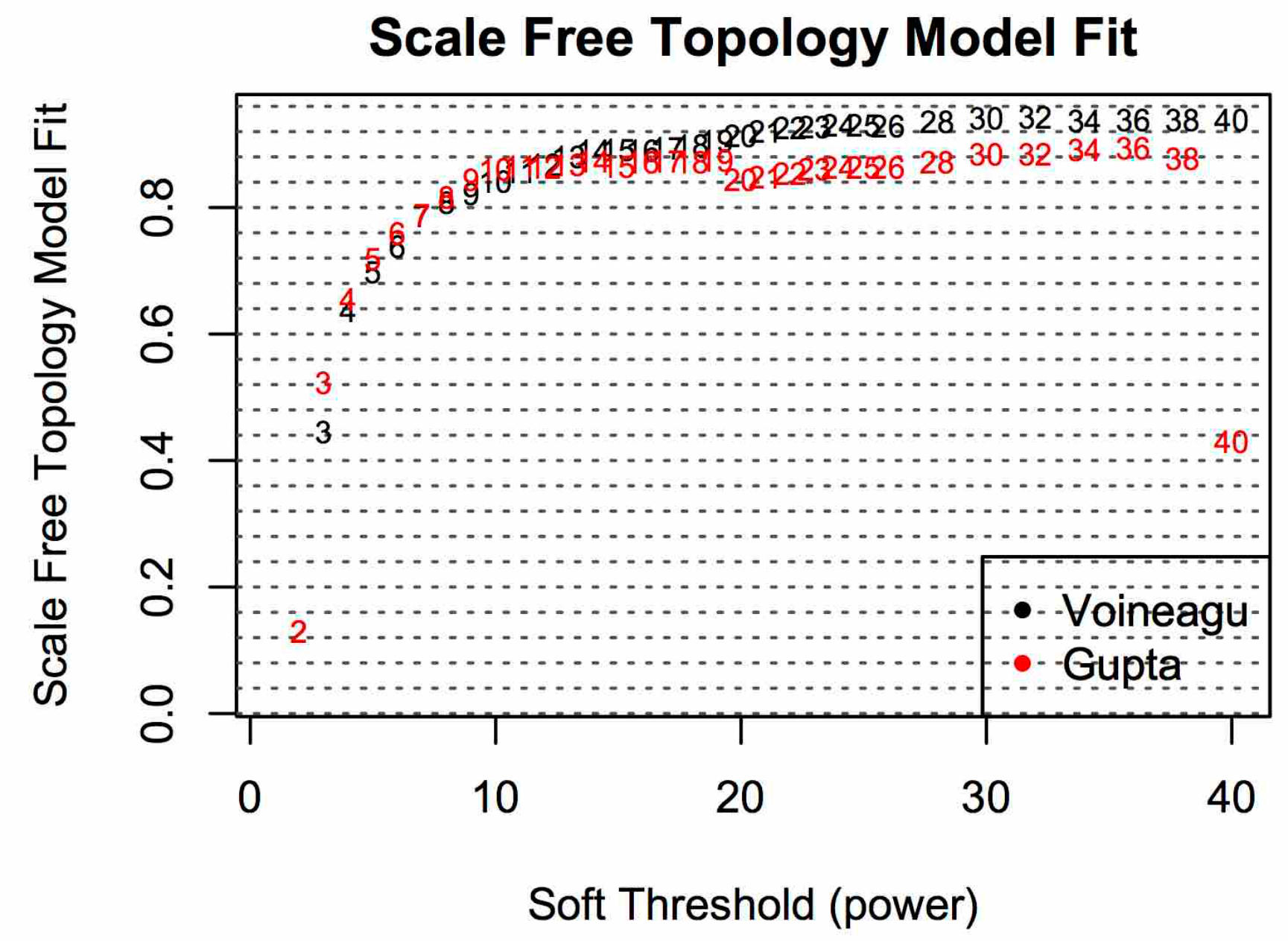
Scale-free topology model fit across a range of soft power thresholds.

This plot shows the scale-free topology model fit scores (R2) across a range of soft power thresholds. This analysis is done in order to choose a soft-power threshold to use in the main analyses. As a rule, we picked the soft power threshold whereby scale-free topology model fit R2 is maximum and begins to plateau (i.e. soft power = 14).

**Fig S2:**
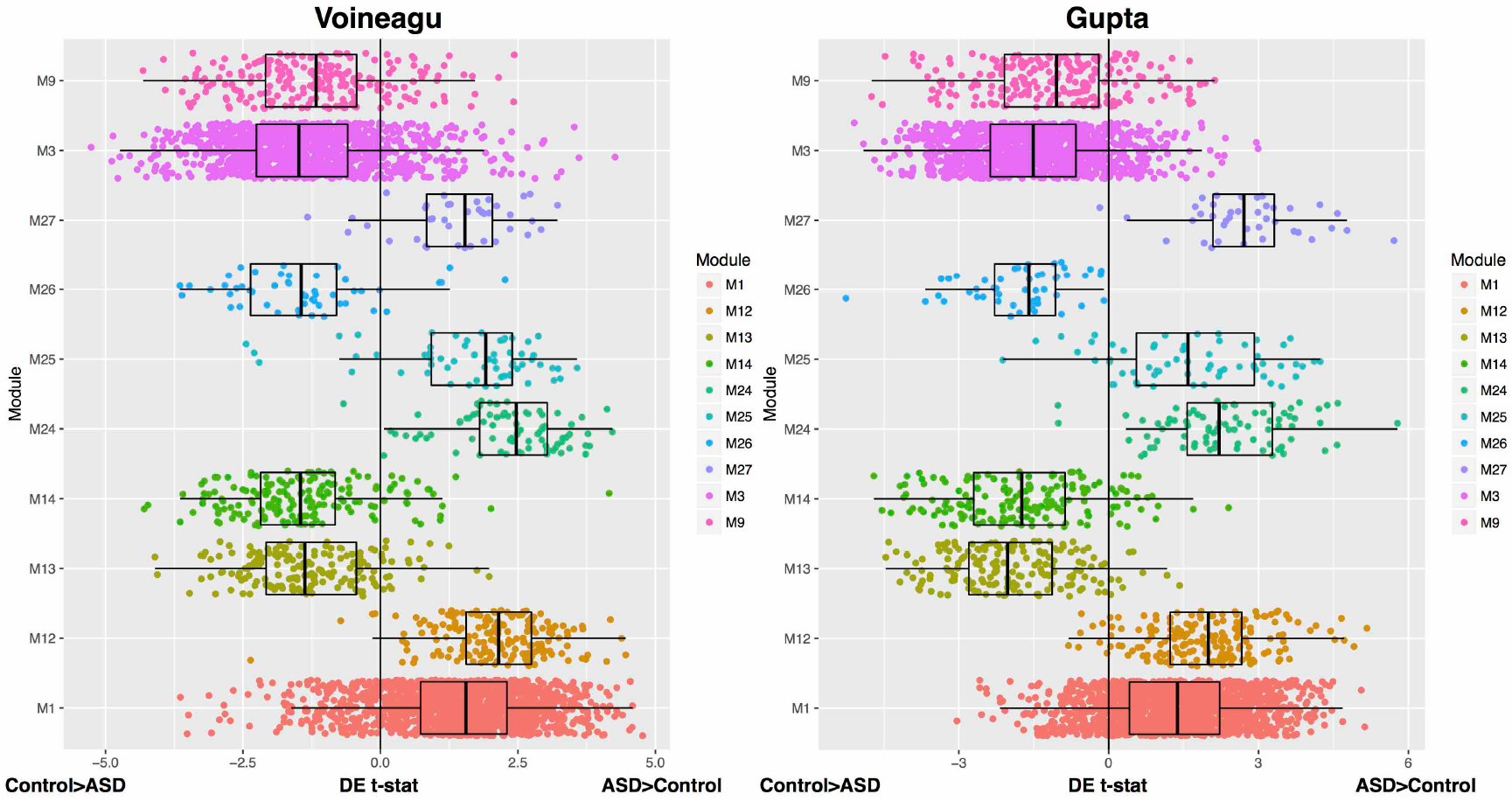
Differential expression load within replicably dysregulated co-expression modules.

This plot shows strength of differential expression (DE) for each gene within the 10 replicably dysregulated co-expression modules. DE strength is quantified continuously as the effect size (t-stat) from the DE gene-level analyses. All modules show a substantial shift in DE signal in the direction congruent with the label of ‘upregulated’ (ASD>Control) or ‘downregulated’ (Control>ASD) given to each module.

**Fig S3:**
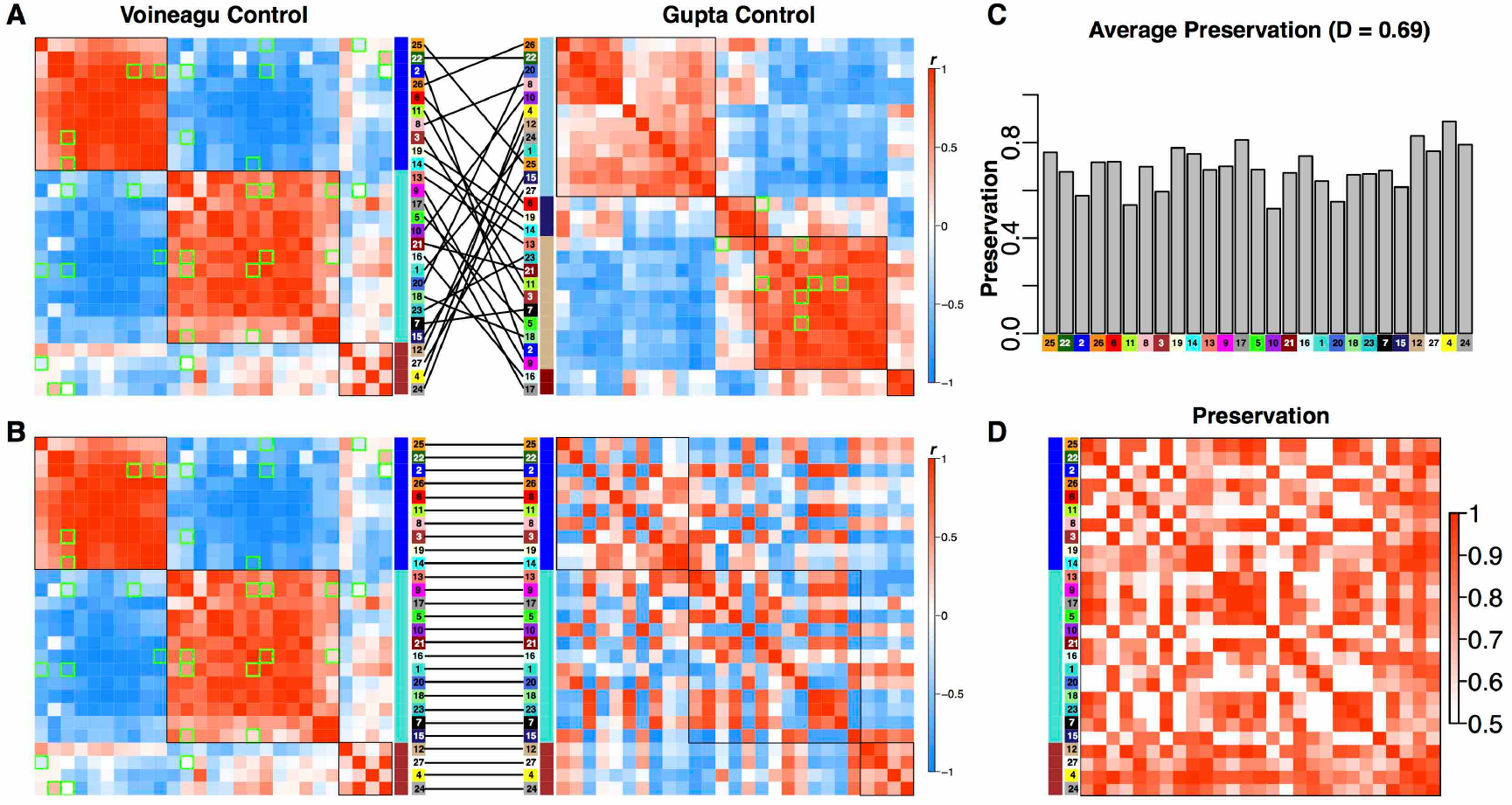
Preservation of eigengene networks in the TD group.

Panel A shows the eigengene networks for Voineagu and Gupta datasets when the rows and columns of the matrix are ordered by meta-module clustering. Panel B shows the matrices when ordered only by the Voineagu TD dataset clustering. Panel C shows average preservation levels across each module. Panel D shows preservation for all pairwise module comparisons. The plots in panels C and D were made using a modified version of the plotEigengeneNetworks function in the WGCNA R library. We modified this function to use ME robust partial correlation matrices.

## Supplemental Table Legends

***Table S1:*** *Enrichments for all dysregulated modules and collections of downregulated and upregulated modules.*

***Table S2:*** *Module membership and hub gene information for each module*

***Table S3:*** *Full result table of analysis examining on-average differential expression in module eigengene variation*

***Table S4:*** *Cell type and cellular component enrichment information*

